# EEG-based classification of natural sounds reveals specialized responses to speech and music

**DOI:** 10.1101/755553

**Authors:** Nathaniel J Zuk, Emily S Teoh, Edmund C Lalor

## Abstract

Humans can easily distinguish many sounds in the environment, but speech and music are uniquely important. Previous studies, mostly using fMRI, have identified separate regions of the brain that respond selectively for speech and music. Yet there is little evidence that brain responses are larger and more temporally precise for human-specific sounds like speech and music, as has been found for responses to species-specific sounds in other animals. We recorded EEG as healthy, adult subjects listened to various types of two-second-long natural sounds. By classifying each sound based on the EEG response, we found that speech, music, and impact sounds were classified better than other natural sounds. But unlike impact sounds, the classification accuracy for speech and music dropped for synthesized sounds that have identical “low-level” acoustic statistics based on a subcortical model, indicating a selectivity for higher-order features in these sounds. Lastly, the trends in average power and phase consistency of the two-second EEG responses to each sound replicated the patterns of speech and music selectivity observed with classification accuracy. Together with the classification results, this suggests that the brain produces temporally individualized responses to speech and music sounds that are stronger than the responses to other natural sounds. In addition to highlighting the importance of speech and music for the human brain, the techniques used here could be a cost-effective and efficient way to study the human brain’s selectivity for speech and music in other populations.

**Highlights:**

- EEG responses are stronger to speech and music than to other natural sounds
- This selectivity was not replicated using stimuli with the same acoustic statistics
- These techniques can be a cost-effective way to study speech and music selectivity

## Introduction

Sound carries a tremendous amount of information about the world around us, and we can easily decipher an acoustic scene. It is clear, though, that speech and music are especially important for humans. They are present in all cultures with a striking amount of diversity (Pinker and Bloom, 1990; Evans and Levinson, 2009; Savage et al., 2015; Youn et al., 2016; Mehr et al., forthcoming). Given their importance, it seems pertinent that our brain should be able to more easily identify these types of sounds relative to other sounds in the natural world. Relatedly, many studies have demonstrated neural specializations to ecologically relevant sounds in other species, particularly to vocalizations and song (King and Nelken, 2009; Theunissen and Elie, 2014). Since speech and music are arguably the most characteristically human of sounds, it is likely that there exist neural specializations in the human brain for them.

Indeed, there is common agreement that specialized regions of the brain responsive to speech and music sounds do exist (Giordano et al., 2014; Hausfeld et al., 2018; Leaver and Rauschecker, 2010; Norman-Haignere et al., 2015; Ogg et al., 2019b; Staeren et al., 2009; Zatorre et al., 2004). These regions appear to be in secondary auditory cortex, which also respond to features beyond those easily captured by spectrotemporal statistics (Kell et al., 2018; Norman-Haignere et al., 2015). However, most of the studies examining specializations for speech and music have used fMRI, and through these studies it is unclear if there are differences in the time course of the responses to speech and music. More recently, an ECoG study showed activation across auditory cortex for speech sounds, and very strong spatially localized neural activation for vocal music in particular (Norman-Haignere et al., 2019). Spatial regions appeared distinct for these activations, but both showed temporally sustained responses throughout the two-second stimulus. A few studies have used EEG to compare neural responses to speech and musical instrument sounds (Cossy et al., 2014; Murray et al., 2006), and to our knowledge only one study has compared time-varying differences in MEG responses to speech and instrument sounds to differences in acoustic features (Ogg et al., 2019a). Still, it is unclear if these responses are unique for speech and music (see Giordano et al., 2014; Norman-Haignere et al., 2015, 2019).

Additionally, there is reason to believe that neural responses to speech and music consistently time-lock to particular acoustic features in these sounds. Several studies have shown greater responses or more consistent patterns of activation to speech sounds than various control sounds where the speech is degraded (Ahissar et al., 2001; Luo and Poeppel, 2007; Nourski et al., 2019; Okada et al., 2010; Peelle et al., 2013; Zoefel et al., 2018) and a similar study found consistent patterns of activation for piano music (Doelling and Poeppel, 2015). Yet auditory cortical neurons can also time-lock to amplitude and frequency modulations in synthetic sounds (deCharms et al., 1998; Liang et al., 2002; Lalor et al., 2009; for review see Joris et al., 2004), and it isn’t clear from previous work if this time-locking is stronger for speech and music than other types of natural sounds.

Here we use classification-based analyses with EEG to identify specialized neural responses to speech and music sounds. Furthermore, unlike other natural sounds that evoke unique temporal responses (such as impact sounds), EEG responds less to sounds that have the same acoustic characteristics as speech and music based on a model of subcortical auditory processing (McDermott and Simoncelli, 2011). Our results demonstrate that the specialization of the brain to speech and music sounds can be observed with EEG, where EEG responses are larger and more time-locked than to other natural or spectrotemporally-matched sounds.

## Material and methods

### Experimental paradigm and EEG preprocessing

This study involved two experiments. In both Experiment 1 and 2, 128 channels of scalp EEG data were recorded, along with two additional channels over the mastoid processes using a Biosemi Active Two system at a sampling rate of 512 Hz. In Experiment 1, six subjects (3 female, ages 24-30) participated. Stimuli consisted of 30 different two-second-long sounds that produced the strongest fMRI responses in the independent components of auditory cortical activity found in a previous study (Norman-Haignere et al., 2015). Of the six speech stimuli used in this experiment, four were unintelligible or in a foreign language, and all six music stimuli were instrumental (see Figure 2 for the list of stimuli). In each trial, the sounds were presented consecutively such that each sound was repeated at least twice during a trial. Five of those sounds were presented a third time immediately following a presentation of the same sound. During the trial, subjects were asked to detect the consecutive repeats (a “one-back” task) by hitting the spacebar. In total, there were 65 sounds per trial, lasting 2 minutes and 10 seconds. Subjects listened to 43-50 trials.

In Experiment 2, 15 subjects participated (7 female, ages 19-31). Stimuli consisted of a subset of five speech and five music sounds from Experiment 1. Additionally, five “impact sounds” were included based on high within-frequency-channel real modulation correlations as determined by a spectrotemporal model of auditory processing (McDermott and Simoncelli, 2011), all of which were sampled from a larger database of natural sounds (Norman-Haignere et al., 2015). The impact sounds were included because they exhibited high classification accuracies in Experiment 1 (see Results below), which were most likely due to the strong, sparse onsets present in these sounds. We also included synthesized (“model-matched”) versions of these 15 sounds that contained identical time-averaged acoustics statistics to the originals based on a subcortical model (McDermott and Simoncelli, 2011). These were included in order to test if the results observed in Experiment 1 were due to some neural selectivity for the spectrotemporal statistics of the sounds. As in Experiment 1, each sound was repeated twice in a trial, but subjects were asked to detect a sound that was identical to the sound before the previous one (a “two-back” task) (**Figure 1a**). This was more difficult than the one-back task used in Experiment 1, and we changed the task in order to make the experiment more engaging. There were 40 trials in total.

**Figure 1:**
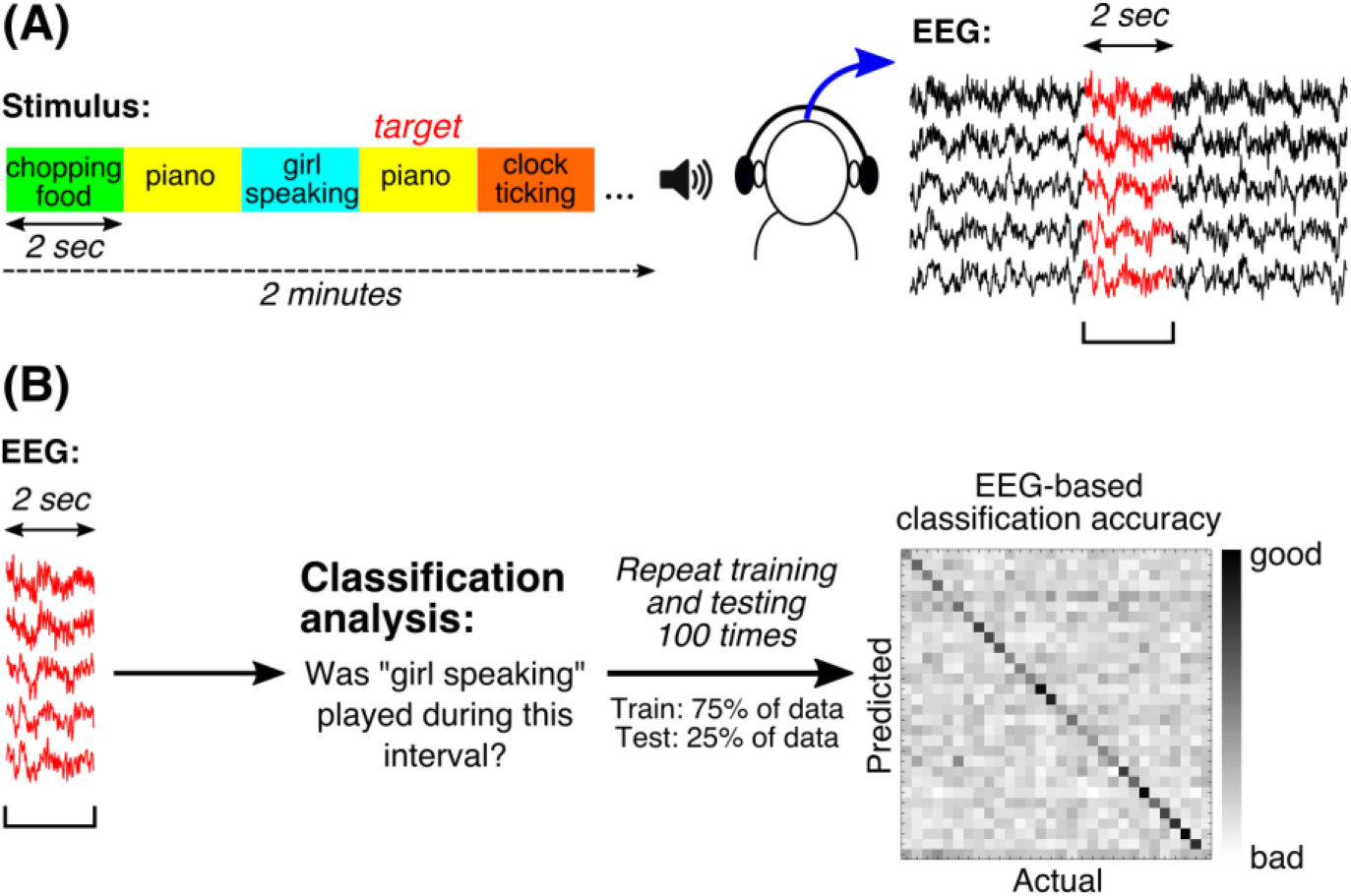
(A) Experiment setup: Stimuli consisted of consecutive two-second sound clips. Subjects were asked to detect repeated sound clips in Experiment 1 (not shown) or detect if a sound clip was identical to the sound clip before the previous one (shown). EEG was recorded and segmented into the corresponding two-second clips after preprocessing. (B) A linear classifier was trained to identify the sound clip presented based on the two-second EEG segments. 75% of the data was used for training and 25% was used for testing. The data was randomly sampled 100 times, and a classification accuracy matrix was constructed based on the average classification accuracy across the 100 repeats of training and testing. For most of the population analyses, the classification accuracies along the diagonal, indicating correct classification, were analyzed.

**Figure 2:**
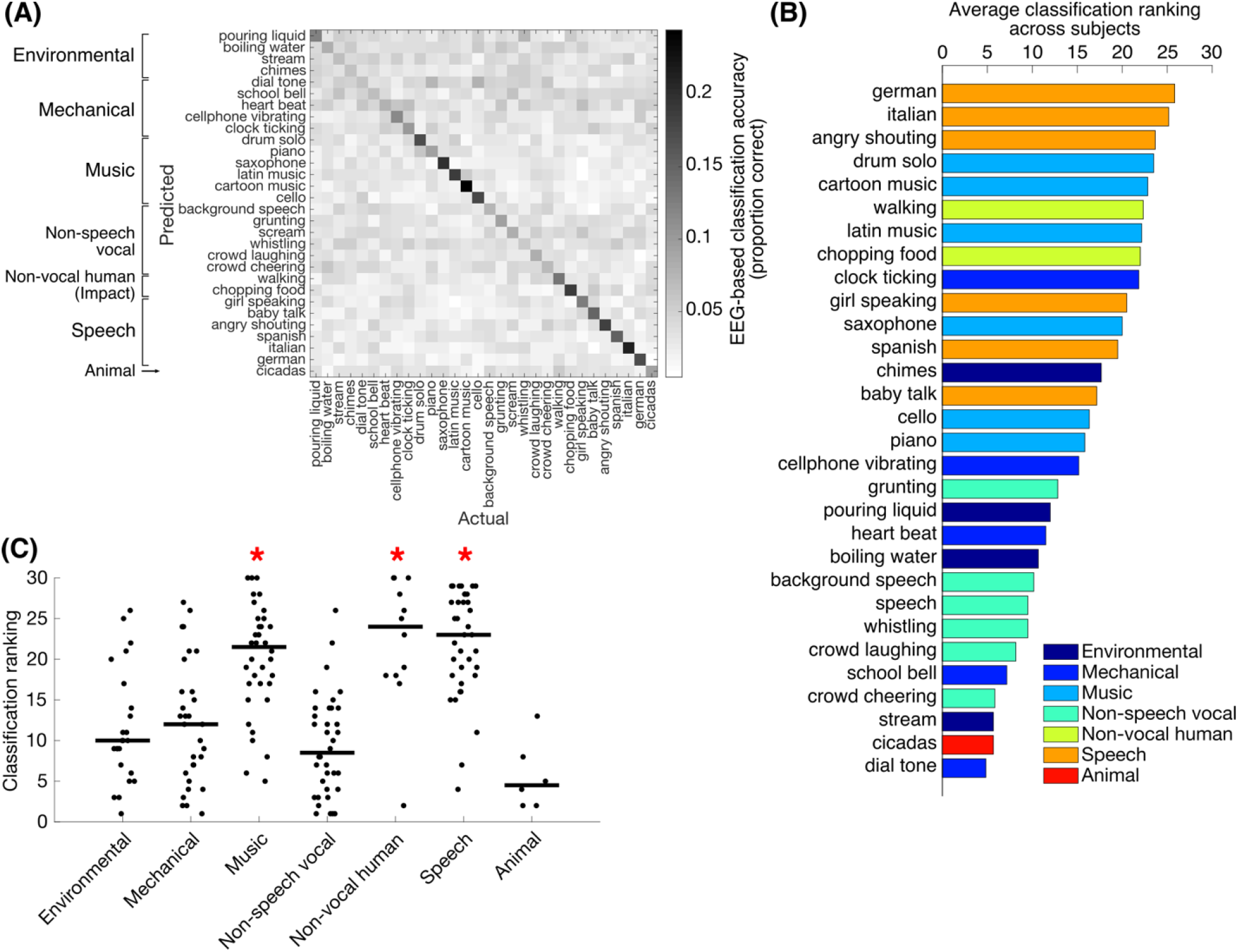
(A) Classification accuracy for an example subject in Experiment 1. None of the stimuli for were classified below chance for this subject (where chance is 0.044 based on the 95% confidence interval from a binomial test with p = 0.033). (B) Ranking of classification accuracy (along the diagonal of the classification matrix in A), averaged across subjects. Music, speech, and “non-vocal human” sounds tend to be classified best. (C) Classification rankings for all types of stimuli across 6 subjects recorded in Experiment 1. Red asterisks indicate that the music, speech, and non-vocal human stimuli were classified significantly better than all other stimuli analyzed (p < 0.05, Dunn & Šidák approach for 21 pairwise comparisons).

### EEG data preprocessing

The 128 scalp channels were referenced to the average of the two mastoids and then filtered between 1-45 Hz using a zero-phase Chebyshev type II filter with a 0.75 to 60 Hz stopband at −60 dB. Eyeblinks and sparse electrical artifacts were identified with the fastICA algorithm (Hyvärinen and Oja, 2000) and removed. Noisy channels were empirically identified based on having a variance that was 3-6 times the interquartile range plus the median across all 128 electrodes and were replaced with a signal equal to the average of the three closest electrodes weighted by the inverse of their distance from the electrode. For one subject with particularly noisy data in Experiment 1, eyeblinks and noisy channels could not be removed reliably using the criteria used for all other subjects, so these preprocessing steps were not performed on this subject and the classification analysis was performed slightly differently for this subject (see below).

### Classification analysis

Each trial of EEG data was spliced into the two-second segments recorded during the sound presentations. Clips that occurred during a target (i.e., repeated) sound were removed from further analysis, resulting in 86-100 EEG clips per stimulus for Experiment 1 and 73-80 EEG clips per stimulus for Experiment 2. In order to reduce the dimensionality of the data, since there were far more dimensions (128 EEG channels × 256 time samples) than data samples for classification, principal components analysis (PCA) was first applied to the spatiotemporal EEG responses for each sound clip and components capturing 95% of the variance of the data (490-1519 components out of 32768 total dimensions) were retained (for the subject with noisy data in Experiment 1, 1049 components capturing 99.999% of the data were retained). The stimulus presented during the two second interval was then identified based on the principal components of the EEG data using multi-class linear discriminant analysis with regularization (*fitcdiscr* in Matlab). 75% of the data were randomly selected for training the classifier and the classifier was tested on the remaining 25% of the data. This selection was repeated 100 times to get 100 classification accuracies for the 30 different stimuli. The classification analysis was done separately for each subject. In order to compare classification accuracies across subjects for population analysis, classification accuracies (along the diagonal on the right in **Figure 1b**) were ranked based on the average classification accuracy across the 100 training-testing repeats.

In order to understand how classification accuracy for the original and model-matched sounds varied over time in Experiment 2, the classification analysis was performed on 200 ms of the EEG response spaced every 100 ms. Because the dimensionality of each 200 ms of EEG data was still larger than the number of samples (128 EEG channels × 26 time samples = 3328 dimensions), PCA was applied to each 200 ms segment of EEG and only the principal components encompassing 95% of the variance of the data were retained for classification (48-496 components). Subsequently, we repeated the first set of classification analyses for both Experiment 1 and 2 using only the first second of the EEG responses, in order to examine how much the classification accuracies change for the speech and music stimuli when the response between 1-2 s is removed. As with the other classification analyses, we used the principal components of the EEG response (319-1143 PCs, 582 components for the subject with noisy data in Experiment 1).

Additionally, to evaluate the contributions of each channel to classification performance, the classification analysis was repeated on each individual EEG channel (256 time samples). To make the procedure comparable to the other analyses, PCA was performed on the data prior to training and testing, but all of the dimensions of the data were retained (256 dimensions).

### Speech/music discrimination analysis

Similar to the classification of individual sounds, we also tested how well sounds could be classified as speech or music based on the EEG responses. For each subject, using only the responses to either speech or music sounds, we retained the principal components that captured 95% of the variance in the responses (290-829 dimensions, and 798 components for the subject with noisy data in Experiment 1 accounting for 99.999% of the variance). We then trained a linear discriminant classifier to identify if a response came from a music sound or a speech sound. Crucially, the classifier was trained on the EEG data from the responses to five of the six speech and music sounds in Experiment 1 and four of the five speech and music sounds in Experiment 2. Then the classifier was tested on the responses to the remaining pair of sounds, resulting in 36 iterations for Experiment 1 and 25 iterations for Experiment 2. This test ensured that the classifier was identifying speech or music sounds based on a general pattern of the EEG response instead of individualized responses to specific sounds.

### Evoked response, global field power (GFP), and phase dissimilarity analyses

In addition to the classification analyses, we examined the median evoked response to each stimulus type across the presentations of non-target stimuli for each subject. As a summary measure of the time course of the evoked response magnitude across the scalp, we also examined the global field power (GFP), which is the standard deviation of the evoked response across channels at each time point (Lehmann and Skrandies, 1980). To evaluate the contribution of response magnitude to the pattern of classification results, for each subject, the GFP was averaged over each two-second interval, and the average GFPs were ranked.

Additionally, we expected that speech and music may evoke temporally consistent responses and, relatedly, consistent response phases over multiple repetitions of the same stimulus, as has been shown by prior work (Doelling and Poeppel, 2015; Luo and Poeppel, 2007). We computed the phase dissimilarity by computing the difference between the phase coherence across trials for a single stimulus and the average phase coherence after sampling trials from all 30 different stimuli. Larger values of phase dissimilarity imply temporally consistent responses for a stimulus from trial to trial. First, phase dissimilarity was computed using 4 Hz frequency bands spaced from 0 to 40 Hz in order to identify the frequency range of the EEG producing the greatest phase consistency across all stimuli and all subjects. Then the phase dissimilarities were ranked for each subject, like the analysis for classification accuracy and average GFP, and differences in phase dissimilarity rankings across were examined across stimulus types.

## Results

### Experiment 1

Nearly all stimuli were classified significantly better than chance for all six subjects; at most 3/30 stimuli failed to reach threshold performance (0.045, which is the 95% criterion for a binomial test with p = 0.033) for each subject (the data for an example subject is shown in **Figure 2a**). However, there appeared to be a considerable amount of variation in the accuracies for correct classification, indicating that some stimuli were easier to classify than others (**Figure 2a**). To quantify the relative magnitude of the classification accuracies for the different types of stimuli, the accuracies for correct classification (along the diagonal of the classification matrix in **Figure 2a**, for example) were ranked. Across the six subjects in Experiment 1, speech, music, and “non-vocal human” sounds were classified significantly better than all other natural sounds (**Figure 2b** **&** **c**; Kruskal-Wallis test: *χ*^2^ = 72.83, *p* < 0.001, pairwise significance in **Figure 2c** was evaluated using t tests with the Dunn & Šidák correction, a modification of Bonferroni correction that is less conservative and more suitable for rankings (Dunn, 1964), for 21 pairwise comparisons, *p* < 0.05).

The “non-vocal human” sounds were two impact sounds characterized by sharp acoustic transients (“chopping food”, “walking on a hard surface”). Using a model that quantified the statistics of the signals within audio frequency and modulation bands based on the physiological stages in subcortical processing (McDermott and Simoncelli, 2011), we identified that these sounds had high within-frequency-channel modulation correlations, for which the real values were notably higher for these two sounds than the other sounds in the dataset. Positive real within-frequency-channel modulation correlations are characteristic of sounds with sharp attacks (McDermott and Simoncelli, 2011). As a result, we hypothesized that these sounds probably evoked transient responses that might have contributed to their increased classification accuracy. In contrast, prior work has shown that speech and music selectivity is sensitive to acoustic features beyond the spectrotemporal statistics captured by this model (Kell et al., 2018; Norman-Haignere et al., 2019, 2015).

### Experiment 2

To validate if the improved classification accuracy for speech, music, and the impact sounds was a result of simpler acoustic statistics, we ran Experiment 2 with speech, music, and impact sounds, as well as model-matched sounds that were generated to have the same frequency and modulation statistics (McDermott and Simoncelli, 2011) (**Figure 3a**). For sounds where the statistics are invariant with time, previous work has shown that model-matched sounds are not discriminable from the originals (McDermott et al., 2013), indicating that the model captures statistics that are sufficient for human perception for some natural sounds such as background sounds (Kell and McDermott, 2019). We then repeated the classification analysis (**Figure 3b**). As in Experiment 1, nearly all stimuli were classified above chance; at most 3/30 stimuli did not pass threshold for each subject. Across all 15 subjects that were recorded during Experiment 2, we found that the model-matched speech and music sounds were classified significantly worse than the originals (Wilcoxon rank-sum test with Bonferroni correction for three comparisons: *p* < 0.001 for both comparisons), while the model-matched impact sounds were classified no differently than the originals (**Figure 3c**). This shows that the neural responses to speech and music producing high classification accuracies were responsive to acoustic features beyond the statistics captured by the model, and therefore cannot be explained by neural specificity to frequency and modulation alone.

**Figure 3:**
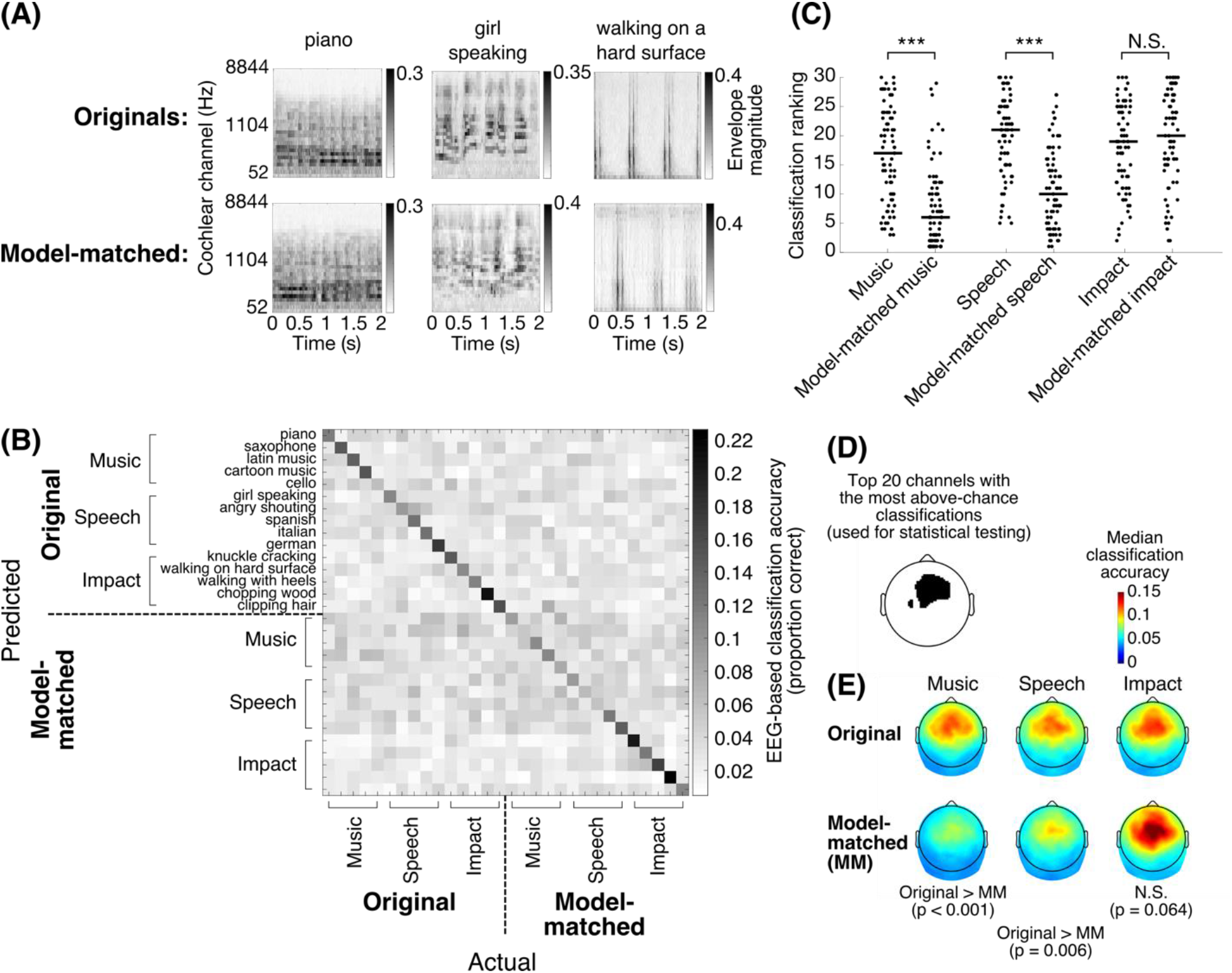
(A) Example cochleograms (see McDermott and Simoncelli, 2011) of the original and model-matched stimulus for an example music (“piano”), speech (“girl speaking”), and impact (“walking on a hard surface”) sound. (B) Classification accuracies for the original and model-matched stimuli for an example subject. All stimuli were classified significantly better than chance (0.047 based on the 95% confidence interval from a binomial test with p = 0.033). The model-matched music and speech had notably worse classification accuracy (along the diagonal) than the originals. (C) Classification rankings for original and model-matched stimuli in Experiment 2 for all 15 subjects. There is a significant reduction in classification ranking for the model-matched music and speech stimuli relative to the originals (Wilcoxon rank-sum with Bonferroni correction for three comparisons: p < 0.001 for both), but no significant difference for the impact sounds. (D) Black designates the top 20 channels that had the most above-chance classifications (p < 0.001 based on a Binomial test) across all stimuli and subjects (377-412 out of 450 classifications). (E) Median classification accuracies across subjects and stimuli for the original and model-matched music, speech, and impact sounds. The significance of the difference between the classification accuracies averaged across the region in (D) is displayed below each column of original and model-matched topographies. Significance was determined with a Wilcoxon rank-sum test with Bonferroni correction for three comparisons. Music and speech show significantly better classifications than their model-matched counterparts, while the impact sounds do not.

Given that previous fMRI work has shown different spatial patterns of selectivity in auditory cortex for speech and music, we also wished to examine the topographies of our EEG classification results for speech and music channel by channel. For all types of the original sounds, classification accuracies were highest in frontal channels. The topography of these classification accuracies may indicate auditory cortical activity (Lalor et al., 2009), but it may also include frontal cortical activity (**Figure 3e**). To test significance of the channel-wise classification accuracies, we focused on 20/128 channels that had the most above-chance classification accuracies for all stimuli and subjects (**Figure 3d**) and compared the classification accuracies averaged across these channels. Both speech and music had significantly better classification accuracies for the original sounds than their model-matched counterparts, while the impact sounds showed no significance (Wilcoxon rank-sum test with Bonferroni correction for three comparisons, significance values are shown in **Figure 3e**).

### Time course of classification accuracy

Next, we repeated the analysis of the EEG data in Experiment 2 using 200 ms blocks of time spaced every 100 ms. This allowed us to examine how classification accuracies and confusion between the stimuli change over the course of the stimulus (**Figure 4a**). Prior work has shown that confusion between images (Cichy et al., 2014; Marti et al., 2015) phonemes (Khalighinejad et al., 2017), and short clips of speech and instrument sounds (Ogg et al., 2019a) varies with time, indicating different stages of processing for the stimuli. However, for the stimuli used in Experiment 2 we found no confusion between classes of stimuli (**Figure 4a**, black line). This could be due to unique temporal responses for the different stimuli.

**Figure 4:**
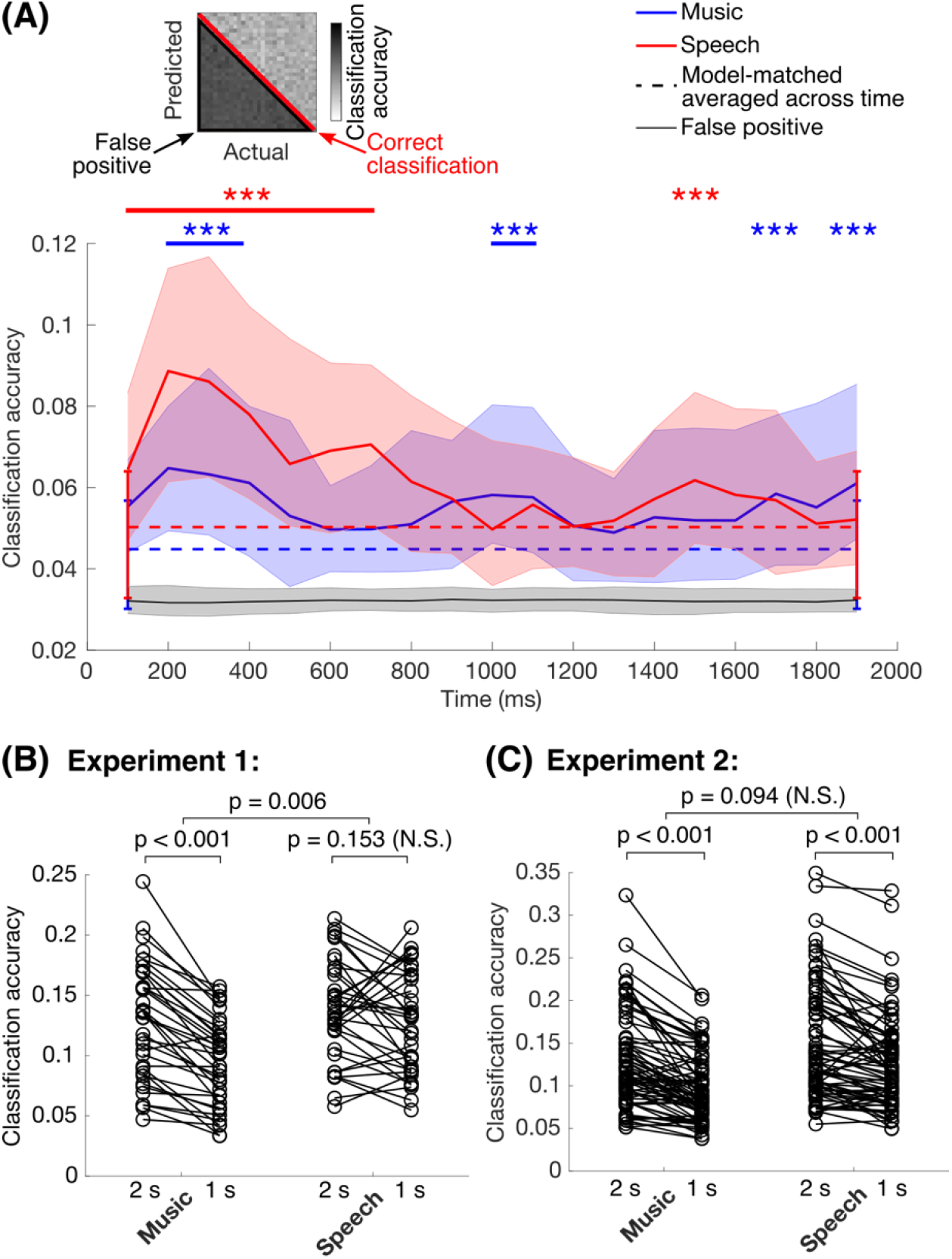
(A) Classification accuracy using 200 ms segments of the EEG response sampled at different points throughout the two-second stimulus was averaged across 15 subjects. We examined the accuracy for correct classification for the different stimulus types (red line), as before, and the off-diagonal classification accuracies indicating the false positive rate (black shaded region). We expected confusion between stimuli, which would result in above-chance false positives. Classification accuracy of the speech (red) and music (blue) stimuli. The solid lines indicate the median values and the shaded regions indicate the interquartile range across all stimuli and subjects. The dashed lines indicate the median and interquartile range of the model-matched stimuli for the speech (red) and music (blue), which were relatively constant over the two second interval. Above, asterisks label 200 ms intervals where the classification accuracy of the original stimuli exceeds the average model matched accuracy (Wilcoxon rank-sum test, FDR with Benjamini-Yekutieli procedure, q < 0.001). The black line and grey region show the median and interquartile range respectively for the false positives. We did not observe confusion between the different stimuli, and the median false positive rate was always slightly but significantly below chance (Wilcoxon signed-rank test relative to chance, with chance probability = 0.033). (C) and (D) show the classification accuracies using the data from Experiment 1 and 2 respectively, when the full two-second response is included (“2 s”) and when only the first second of the response is used (“1 s”). Comparisons between 2 s and 1 s use Wilcoxon signed-rank test, and comparisons of those differences between stimuli use the rank-sum test. Classification accuracy tend to decrease for all stimuli when only the first second of the response is used (this does not reach significance for speech in Experiment 1), but the change is a slightly larger change in accuracy for music than for speech. The difference in accuracy reaches significance for Experiment 1, but it doesn’t reach significance for Experiment 2.

We then compared the classification accuracies for the original speech and music to the average classification accuracies of the model-matched versions. Classification of speech was significantly better than its model-matched counterpart mainly between 100-700 ms after stimulus onset, and the classification at 1150 ms was also significant (**Figure 4a**). Music similarly showed significantly better classification within the early time range (specifically 200-400 ms), but it also continued to show significantly better classification 1000-1100 ms, 1700 ms, and 1900 ms after stimulus onset.

The time points exhibiting significance beyond one second were short, and it was not entirely clear how important these later response times are for representing the stimulus. To validate this, we repeated the classification analysis in Experiments 1 and 2 using only the first second of the response to the stimuli. We expected that the classification accuracy would decrease for both music and speech stimuli when the last second of the response was not included simply because of the reduction of information available for classification, but we hypothesized that this decrease would be larger for music than for speech if the time points after 1000 ms were important. We found a weak but significantly larger decrease for music than for speech in Experiment 1 (**Figure 4b**), and we found a similar pattern in the data for Experiment 2 that did not reach significance (**Figure 4c**). Thus, the response after 1000 ms may be carrying more information for music than for speech, but the contribution appears to be very weak.

### EEG-based discrimination between speech and music

Using both sets of data from Experiments 1 and 2, we created classifiers that identified whether a sound clip was speech or music based on the EEG response. Importantly, the classifier was tested on EEG responses for stimuli that were not included in training. This was to ensure that the classifier was not relying on stimulus-specific temporal responses and was instead relying a more general spatiotemporal response to speech or music. For both sets of data, speech/music discrimination was not as successful as classification of individual sounds: for Experiment 1 speech and music were not classified above chance for any of the subjects, and for Experiment 2 either speech or music exhibited above chance performance for only 2/15 subjects (above chance criterion was 0.56 for the 95% confidence interval based on a binomial test with p=0.5). This supports our expectation that the responses are temporally individualized for each sound. Additionally, it suggests that EEG does not pick up the subtle spatial differences in cortical responsiveness to speech and music sounds, as has been found in prior fMRI literature (Giordano et al., 2014; Norman-Haignere et al., 2019, 2015; Ogg et al., 2019b) and with MEG (Ogg et al., 2019a), which might be observed with many more speech and music samples in a future experiment.

### Evoked response, GFP, and phase dissimilarity analyses

One possible explanation for our classification results is that music and speech elicit larger responses than other natural or model-matched sounds, because if the responses are larger, they would be easier to classify based on the EEG. To evaluate this hypothesis, we did a more standard analysis of the evoked responses to each of the different sounds: We looked at the evoked EEG response and the time-varying GFP for each of the different stimulus types (**Figure 5a** **&** **b**). Typically, these average responses would be used to identify differences in the neural responses to the different types of sounds, presuming that the responses have a consistent time course within each class of stimuli. There is a clear frontal positivity in the response for all of the different classes of sounds (**Figure 5a** **&** **b**), but there were no obvious differences in the time course for these different stimuli. This is expected if the stimulus-specific response was temporally individualized, as we found in our time-based analysis (**Figure 4a**) and our speech/music discrimination analysis.

**Figure 5:**
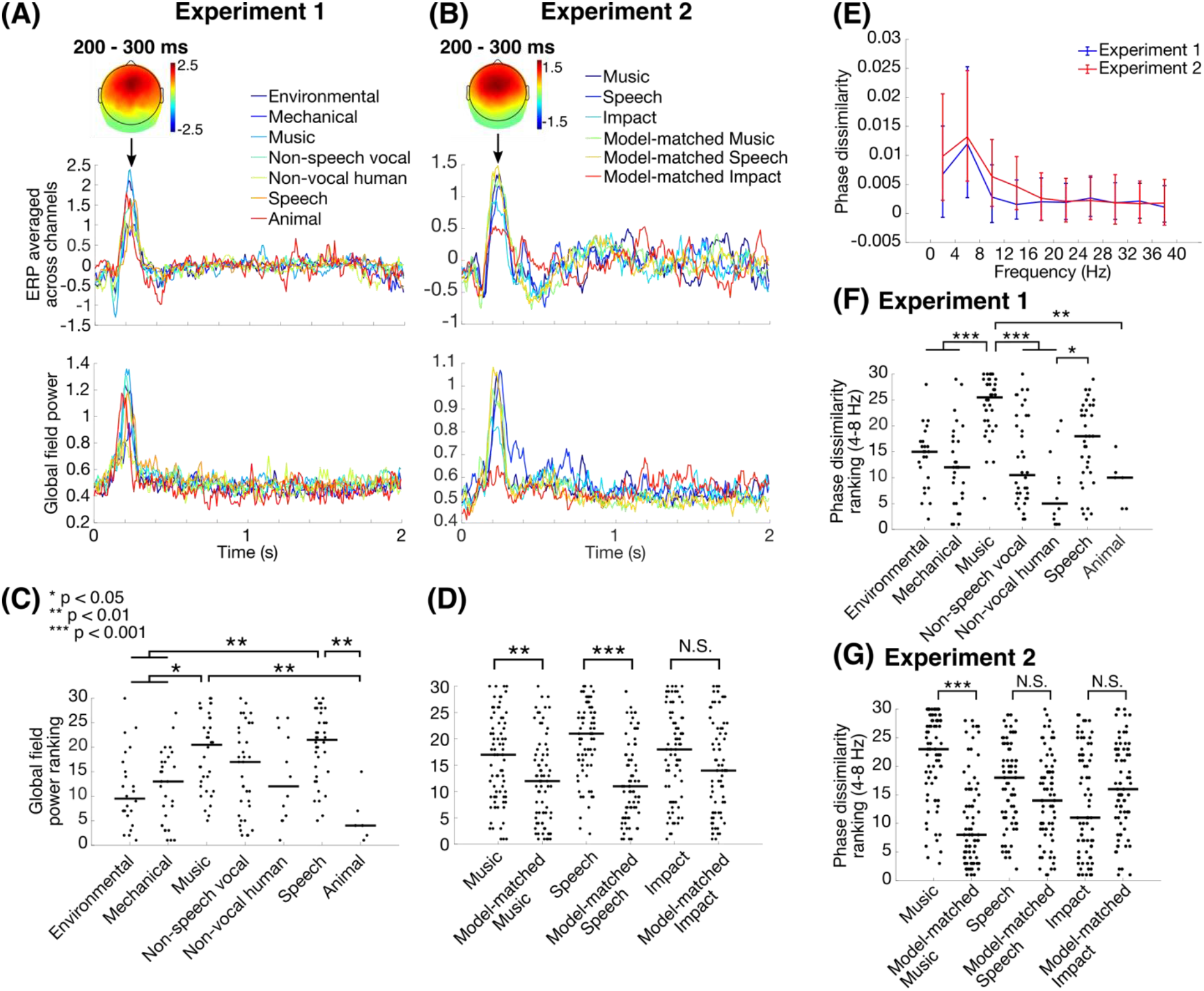
(A) and (B) show the evoked responses for Experiments 1 and 2 respectively. Top plots show the evoked response averaged across all EEG electrodes, where each line is the median electrode-averaged evoked response across stimuli and subjects; for clarity error bars are not shown. Bottom plots show the global field power for each stimulus type, where each line is the median across stimuli and subjects. The topographies of the evoked response between 200 – 300 ms, median values across all stimuli and subjects, are shown at the top. For each stimulus, the global field power of the evoked response was averaged over the two-second duration of the stimulus, and the average global field powers for each of the stimuli were ranked. (C) and (D) are these GFP rankings for all subjects and stimuli for Experiments 1 and 2, respectively. Despite the lack of clear temporal response differences between the evoked responses for the different stimuli, the pattern of GFP rankings are similar to those we observed based on classification accuracy, indicating that the classification accuracy is partly driven by subtle differences in the magnitude of the neural responses to these sounds. (C: Kruskal-Wallis test: *χ*^2^ = 36.15, *p* < 0.001, asterisks based on pairwise t-test with Dunn & Šidák correction for 21 comparisons; D: significance based on Wilcoxon rank-sum test with Bonferroni correction for the three comparisons shown). (E) Phase dissimilarity as a function of frequency for Experiment 1 (blue) and Experiment 2 (red), computed for each 4 Hz frequency band between 0 and 40 Hz. Lines show median values across all stimuli and subjects, and error bars show the interquartile range. On average, phase dissimilarity was highest between 4-8 Hz. (F & G) Ranked phase dissimilarity (Luo and Poeppel, 2007) for all stimuli and subjects in Experiment 1 and Experiment 2. To rank the phase dissimilarity, we focused on the 4-8 Hz range because it produced the largest average phase dissimilarity for all stimuli and subjects. Below are the phase dissimilarity rankings for Experiment 1 (F) and Experiment 2 (G). Phase dissimilarity rankings generally showed the same pattern as classification accuracies and average GFP, but speech showed phase dissimilarity that was not significantly greater than most of the other natural sounds nor its model-matched counterpart, while the significant differences relative to music were largely retained (F: Kruskal-Wallis test: *χ*^2^ = 56.55, *p* < 0.001; asterisks in F & G use the same statistical tests as those used in C and D).

After computing the evoked responses, we also averaged the GFP over the two-second interval of each response. The average GFP values were then ranked within each subject in order to compare them across subjects, like we did for the classification accuracies. Surprisingly, the pattern of average GFP rankings mimicked the pattern of classification accuracy rankings for both experiments, albeit with slightly weaker differences between stimulus types (**Figure 5c** **&** **d**, compare to Figure 2c and **Figure 3c** for Experiments 1 and 2 respectively). This indicates that the differences in classification accuracy result partly from differences in the magnitude of the brain’s response to the different sounds. However, the differences in response magnitudes are subtle, encompassing stimulus-specific responses throughout the full two-second interval of the stimulus, which could not be easily seen by looking at the averaged time course of the GFP.

Prior work has shown that neural responses for both speech and music have consistent low-frequency phases (below 8 Hz) indicating consistent temporal responses across repetitions of the same stimulus (Doelling and Poeppel, 2015; Luo and Poeppel, 2007). We quantified this phase consistency using phase dissimilarity, which is the difference between the within-stimulus phase coherence and the between-stimulus phase coherence (Luo and Poeppel, 2007). We first computed the phase dissimilarity for 4 Hz frequency bands and found that for both experiments the median phase dissimilarity across all stimuli and subjects was largest between 4-8 Hz. The phase dissimilarities within this frequency band were then ranked. Like the result using average GFP, the phase dissimilarity generally produced a similar pattern of rankings as the classification accuracies for both experiments, but interestingly the difference between the phase dissimilarity rankings for the speech and model-matched speech stimuli was not significant (**Figure 5f** **&** **g**). We also validated that the median phase dissimilarity was largest for the speech and music stimuli in both experiments, so the differences in phase dissimilarity rankings we observed were not a consequence of the choice of frequency band. Thus, neural responses between 4-8 Hz appear to be more time-locked to stimulus repetitions than other natural or model-matched stimuli, particularly for music.

## Discussion

In this study, we showed that EEG-based classification performs better for speech and music sounds than for other natural sounds. Additionally, unlike the responses to impact sounds, this responsiveness did not persist for model-matched stimuli generated to have statistics that are known to be captured by subcortical processing (McDermott and Simoncelli, 2011). The pattern persisted, albeit more weakly, when we examined the average GFP for each of the sounds and their phase dissimilarity between 4-8 Hz. Together, these results indicate that the human brain is especially responsive to naturalistic speech and music sounds.

To evaluate the effect on our results of basic acoustic features that are known to be processed subcortically and in primary auditory cortical areas, we used a model that captures the statistics of the acoustics accounted for by these processing stages. This model was then used to resynthesize “model-matched” sounds with identical statistics for these features, namely the frequency spectrum, modulation spectrum, and correlations between different frequency and modulation bands (see also Kell et al., 2018; Norman-Haignere et al., 2015; Norman-Haignere and McDermott, 2018). Some of the acoustic complexities lost in this model include pitch and rhythmic structure (McDermott and Simoncelli, 2011), for which the biophysical mechanisms involved in capturing these attributes are less clear. While it is known that pitch and rhythmic structure are important for representing speech and music, our main interest was in evaluating if the differences observed here can be explained by known biophysical acoustic processing stages that are not unique for speech and music. Pinpointing exactly which mechanisms are involved in producing neural selectivity to these other acoustic complexities will require further work.

We also showed that there are differences in the timing of neural responses for speech and music, such that classification accuracy peaked 200 ms after onset for speech but persisted sporadically throughout the stimulus for music. Furthermore, we found that the classification accuracy decreased more for music than for speech when the response from 1000-2000 ms was left out, although this effect was very weak. Both of these results are in line with recent ECoG work (Norman-Haignere et al., 2019). In particular, the component responsive to music gradually increased over the two-second duration of the stimulus, peaking just after one second (Norman-Haignere et al., 2019). Contrary to our expectations, though, we found no confusion between the sounds over time, and we were also largely unable to discriminate between speech and music sounds based on the EEG responses in either experiment. Relatedly, the short, above-model-matched classification accuracies beyond 1000 ms that we observed for music could in part be due to features in the specific stimuli we chose; this could be validated in future studies using other music stimuli. But while differences in temporal processing of speech and music generally may be weakly present in EEG, our results based on the lack of confusion between stimuli and based on phase dissimilarity more prominently demonstrate that each response is aligned to temporal features that are unique for individual speech and music sounds. However, the temporal features that generate these responses may differ. Midbrain neuron models with different synaptic parameters optimally capture vowel formant and beat-related information in speech and music respectively (Carney et al., 2015; Zuk et al., 2018). The specializations for speech and music in the auditory cortex may have underpinnings at earlier processing stages in the auditory system.

Much of our focus was on the speech and music results, but the importance of decoding impact sounds also reflects several earlier studies. In particular, the impact sounds used in our study could all be categorized as “non-human” or “non-living” using terms from prior work (Cossy et al., 2014; Giordano et al., 2013; Lewis et al., 2005; Murray et al., 2006). There may also be a particular processing stream for these types of stimuli as well; not only are these sounds decodable with fMRI (Hjortkjær et al., 2018), but these types of sounds have been shown to evoke motor cortical areas (Lewis et al., 2005). In the absence of any clear evidence of this in the present work, a parsimonious interpretation would be to assume that, in Experiments 1 and 2, the impact sounds were well classified because of differences in the timing of transient events that produced time-locked evoked potentials.

The spatial organization of EEG scalp electrodes that were involved in decoding these sounds tended to be frontal electrodes. The topography generated by this pattern of classification accuracies is indicative of auditory cortical activity (Lalor et al., 2009). However, we cannot completely rule out the possibility that frontal regions of cortex are involved. Frontal cortex is known to be involved in task performance (Gold and Shadlen, 2007; Miller and Cohen, 2001) and neurons in frontal cortex show a specificity for evoked sounds that are task relevant (Fritz et al., 2010). In our experiments, subjects were required to respond to repeated sounds in our experiment, so the task did require some amount of mental categorization of the sounds, which could have evoked frontal cortical regions, even for non-target sounds. One possible way of reducing the task-related activity, for example, is by presenting the sounds during a completely unrelated visual task. Even then, there is the possibility that frontal cortical neurons are already tuned to ecologically relevant sounds as a result of general experience. All of the natural sounds were recognizable and identifiable (see the behavioral study from Norman-Haignere et al., 2015). Thus, it would be very difficult to completely isolate or rule out the possibility of frontal activity involvement using EEG, and well beyond the scope of this study.

Given that midbrain-level processing of basic acoustic features and frontal cortex selectivity could both be involved in generating speech and music selectivity in auditory cortical regions, it is still an open question as to whether the selectivity we observe here is a result of “bottom-up” processing, due to auditory processing that does not require subject engagement, or “top-down” factors that depend upon the subject’s cognitive state. Disambiguating these two is tricky because specific stimuli with particular acoustic characteristics such as sparsity and roughness could engage a subject’s attention (Huang and Elhilali, 2017; Zhao et al., 2019). However, we think it is unlikely that subjects were actively attending to speech and music more than other sounds in the stimulus set because the behavioral task places no special emphasis on these stimuli and all stimuli likely became less interesting over many repeats (>80 presentations). Understanding the exact effects of attention on the speech and music selectivity we have observed here will require further work.

Our work and that of others have identified neural responses specialized for processing music. Yet the definition of music is still debatable. In the field of ethnomusicology the definition of the ability to create music is more often used because of the ambiguity of defining music (Miller and Shahriari, 2012; Rice, 2014). In spite of this others have shown that there is general consensus in identifying “musical” sounds amidst a collection of commonly heard sound recordings, including the ones used in our study (see Norman-Haignere et al., 2015). Our paradigm provides an opportunity to study what features make the music-selective regions of the brain respond and how this specialization is affected by culture, experience, and subject expectations (Di Liberto et al., 2019).

## Conclusions

In this study, we showed that EEG responses to speech and music are stronger and more time-locked than to other natural and model-matched sounds. EEG is cheaper and more portable than fMRI, and even though it lacks spatial resolution, we showed that we could still detect specializations for speech and music. With clever experimental design, this opens up possibilities to study the neural processing of music where fMRI is less feasible, such as with infants or populations in remote geographic locations.

## Acknowledgments

The authors would like to thank Sam Norman-Haignere and Josh McDermott for providing the stimuli of natural sounds used in our experiments, and Lauren Szymula for collecting EEG data in Experiment 2.

## References

Ahissar, E., Nagarajan, S., Ahissar, M., Protopapas, A., Mahncke, H., Merzenich, M.M., 2001. Speech comprehension is correlated with temporal response patterns recorded from auditory cortex. Proc. Natl. Acad. Sci. 98, 13367–13372. https://doi.org/10.1073/pnas.201400998

Carney, L.H., Li, T., McDonough, J.M., 2015. Speech coding in the brain: representation of vowel formants by midbrain neurons tuned to sound fluctuations. eneuro 2. https://doi.org/10.1523/ENEURO.0004-15.2015

Cichy, R.M., Pantazis, D., Oliva, A., 2014. Resolving human object recognition in space and time. Nat. Neurosci. 17, 455–462. https://doi.org/10.1038/nn.3635

Cossy, N., Tzovara, A., Simonin, A., Rossetti, A.O., De Lucia, M., 2014. Robust discrimination between EEG responses to categories of environmental sounds in early coma. Front. Psychol. 5, 155. https://doi.org/10.3389/fpsyg.2014.00155

deCharms, R.C., Blake, D.T., Merzenich, M.M., 1998. Optimizing sound features for cortical neurons. Science 280, 1439–1443. https://doi.org/10.1126/science.280.5368.1439

Di Liberto, G.M., Pelofi, C., Bianco, R., Patel, P., Mehta, A.D., Herrero, J.L., Cheveigné, A. de, Shamma, S., Mesgarani, N., 2019. Cortical encoding of melodic expectations in human temporal cortex. bioRxiv 714634. https://doi.org/10.1101/714634

Doelling, K.B., Poeppel, D., 2015. Cortical entrainment to music and its modulation by expertise. Proc. Natl. Acad. Sci. 112, E6233–E6242. https://doi.org/10.1073/pnas.1508431112

Dunn, O.J., 1964. Multiple comparisons using rank sums. Technometrics 6, 241–252. https://doi.org/10.1080/00401706.1964.10490181

Evans, N., Levinson, S.C., 2009. The myth of language universals: language diversity and its importance for cognitive science. Behav. Brain Sci. 32, 429–448. https://doi.org/10.1017/S0140525X0999094X

Fritz, J.B., David, S. V, Radtke-Schuller, S., Yin, P., Shamma, S.A., 2010. Adaptive, behaviorally gated, persistent encoding of task-relevant auditory information in ferret frontal cortex. Nat. Neurosci. 13, 1011–1019. https://doi.org/10.1038/nn.2598

Giordano, B.L., McAdams, S., Zatorre, R.J., Kriegeskorte, N., Belin, P., 2013. Abstract encoding of auditory objects in cortical activity patterns. Cereb. Cortex 23, 2025–2037. https://doi.org/10.1093/cercor/bhs162

Giordano, B.L., Pernet, C., Charest, I., Belizaire, G., Zatorre, R.J., Belin, P., 2014. Automatic domain-general processing of sound source identity in the left posterior middle frontal gyrus. Cortex 58, 170–185. https://doi.org/10.1016/J.CORTEX.2014.06.005

Gold, J.I., Shadlen, M.N., 2007. The neural basis of decision making. Annu. Rev. Neurosci. 30, 535–574. https://doi.org/10.1146/annurev.neuro.29.051605.113038

Hausfeld, L., Riecke, L., Formisano, E., 2018. Acoustic and higher-level representations of naturalistic auditory scenes in human auditory and frontal cortex. Neuroimage 173, 472–483. https://doi.org/10.1016/J.NEUROIMAGE.2018.02.065

Hjortkjær, J., Kassuba, T., Madsen, K.H., Skov, M., Siebner, H.R., 2018. Task-modulated cortical representations of natural sound source categories. Cereb. Cortex 28, 295–306. https://doi.org/10.1093/cercor/bhx263

Huang, N., Elhilali, M., 2017. Auditory salience using natural soundscapes. J. Acoust. Soc. Am. 141, 2163–2176. https://doi.org/10.1121/1.4979055

Hyvärinen, A., Oja, E., 2000. Independent component analysis: algorithms and applications. Neural Networks 13, 411–430. https://doi.org/10.1016/S0893-6080(00)00026-5

Joris, P.X., Schreiner, C.E., Rees, A., 2004. Neural processing of amplitude-modulated sounds.Physiol. Rev. 84, 541–577. https://doi.org/10.1152/physrev.00029.2003

Kell, A.J.E., McDermott, J.H., 2019. Invariance to background noise as a signature of non-primary auditory cortex. Nat. Commun. 10, 3958. https://doi.org/10.1038/s41467-019-11710-y

Kell, A.J.E., Yamins, D.L.K., Shook, E.N., Norman-Haignere, S. V., McDermott, J.H., 2018. A Task-optimized neural network replicates human auditory behavior, predicts brain responses, and reveals a cortical processing hierarchy. Neuron 98, 630–644. https://doi.org/10.1016/J.NEURON.2018.03.044

Khalighinejad, B., Cruzatto da Silva, G., Mesgarani, N., 2017. Dynamic encoding of acoustic features in neural responses to continuous speech. J. Neurosci. 37, 2176–2185. https://doi.org/10.1523/JNEUROSCI.2383-16.2017

King, A.J., Nelken, I., 2009. Unraveling the principles of auditory cortical processing: can we learn from the visual system? Nat. Neurosci. 12, 698–701. https://doi.org/10.1038/nn.2308

Lalor, E.C., Power, A.J., Reilly, R.B., Foxe, J.J., 2009. Resolving precise temporal processing properties of the auditory system using continuous stimuli. J. Neurophysiol. 102, 349–359. https://doi.org/10.1152/jn.90896.2008

Leaver, A.M., Rauschecker, J.P., 2010. Cortical representation of natural complex sounds: effects of acoustic features and auditory object category. J. Neurosci. 30, 7604–7612. https://doi.org/10.1523/jneurosci.0296-10.2010

Lehmann, D., Skrandies, W., 1980. Reference-free identification of components of checkerboard-evoked multichannel potential fields. Electroencephalogr. Clin. Neurophysiol. 48, 609–621. https://doi.org/10.1016/0013-4694(80)90419-8

Lewis, J.W., Brefczynski, J.A., Phinney, R.E., Janik, J.J., DeYoe, E.A., 2005. Distinct cortical pathways for processing tool versus animal sounds. J. Neurosci. 25, 5148–5158. https://doi.org/10.1523/jneurosci.0419-05.2005

Liang, L., Lu, T., Wang, X., 2002. Neural representations of sinusoidal amplitude and frequency modulations in the primary auditory cortex of awake primates. J Neurophysiol 87, 2237–2261. https://doi.org/10.1152/jn.2002.87.5.2237

Luo, H., Poeppel, D., 2007. Phase patterns of neuronal responses reliably discriminate speech in human auditory cortex. Neuron 54, 1001–1010. https://doi.org/10.1016/j.neuron.2007.06.004

Marti, S., King, J.-R., Dehaene, S., 2015. Time-resolved decoding of two processing chains during dual-task interference. Neuron 88, 1297–1307. https://doi.org/10.1016/J.NEURON.2015.10.040

McDermott, J.H., Schemitsch, M., Simoncelli, E.P., 2013. Summary statistics in auditory perception. Nat Neurosci 16, 493–498. https://doi.org/10.1038/nn.3347

McDermott, J.H., Simoncelli, E.P., 2011. Sound texture perception via statistics of the auditory periphery: evidence from sound synthesis. Neuron 71, 926–940. https://doi.org/10.1016/j.neuron.2011.06.032

Mehr, S., Singh, M., Knox, D., Ketter, D., Pickens-Jones, D., Atwood, S., Lucas, C., Egner, A., Jacoby, N., Hopkins, E.J., Howard, R.M., O’Donnell, T.J., Pinker, S., Krasnow, M., Glowacki, L., (forthcoming). Universality and diversity in human song. Science. https://doi.org/10.31234/OSF.IO/EMQ8R

Miller, E.K., Cohen, J.D., 2001. An integrative theory of prefrontal cortex function. Annu. Rev. Neurosci. 24, 167–202. https://doi.org/10.1146/annurev.neuro.24.1.167

Miller, T.E., Shahriari, A., 2012. World Music: A Global Journey, Third Edition. ed. Routledge, New York, NY. https://doi.org/10.4324/9780203892169

Murray, M.M., Camen, C., Gonzalez Andino, S.L., Bovet, P., Clarke, S., 2006. Rapid brain discrimination of sounds of objects. J. Neurosci. 26, 1293–302. https://doi.org/10.1523/JNEUROSCI.4511-05.2006

Norman-Haignere, S., Feather, J., Brunner, P., Ritaccio, A., McDermott, J.H., Schalk, G., Kanwisher, N., 2019. Intracranial recordings from human auditory cortex reveal a neural population selective for musical song. bioRxiv 696161. https://doi.org/10.1101/696161

Norman-Haignere, S., Kanwisher, N.G., McDermott, J.H., 2015. Distinct cortical pathways for music and speech revealed by hypothesis-free voxel decomposition. Neuron 88, 1281–1296. https://doi.org/10.1016/j.neuron.2015.11.035

Norman-Haignere, S. V., McDermott, J.H., 2018. Neural responses to natural and model-matched stimuli reveal distinct computations in primary and nonprimary auditory cortex. PLOS Biol. 16, e2005127. https://doi.org/10.1371/journal.pbio.2005127

Nourski, K. V., Steinschneider, M., Rhone, A.E., Kovach, C.K., Kawasaki, H., Howard, M.A., 2019. Differential responses to spectrally degraded speech within human auditory cortex: An intracranial electrophysiology study. Hear. Res. 371, 53–65. https://doi.org/10.1016/J.HEARES.2018.11.009

Ogg, M., Carlson, T.A., Slevc, R., 2019a. The rapid emergence of auditory object representations in cortex reflect central acoustic attributes. J. Cogn. Neurosci. 1–13. https://doi.org/10.1162/jocn_a_01472

Ogg, M., Moraczewski, D., Kuchinsky, S.E., Slevc, L.R., 2019b. Separable neural representations of sound sources: speaker identity and musical timbre. Neuroimage 191, 116–126. https://doi.org/10.1016/J.NEUROIMAGE.2019.01.075

Okada, K., Rong, F., Venezia, J., Matchin, W., Hsieh, I.-H., Saberi, K., Serences, J.T., Hickok, G., 2010. Hierarchical organization of human auditory cortex: evidence from acoustic invariance in the response to intelligible speech. Cereb. Cortex 20, 2486–2495. https://doi.org/10.1093/cercor/bhp318

Peelle, J.E., Gross, J., Davis, M.H., 2013. Phase-locked responses to speech in human auditory cortex are enhanced during comprehension. Cereb. Cortex 23, 1378–1387. https://doi.org/10.1093/cercor/bhs118

Pinker, S., Bloom, P., 1990. Natural language and natural selection. Behav. Brain Sci. 13, 707–727. https://doi.org/10.1017/S0140525X00081061

Rice, T., 2013. Ethnomusicology: A Very Short Introduction. Oxford University Press.

Savage, P.E., Brown, S., Sakai, E., Currie, T.E., 2015. Statistical universals reveal the structures and functions of human music. Proc. Natl. Acad. Sci. U. S. A. 112, 8987–8992. https://doi.org/10.1073/pnas.1414495112

Staeren, N., Renvall, H., De Martino, F., Goebel, R., Formisano, E., 2009. Sound categories are represented as distributed patterns in the human auditory cortex. Curr. Biol. 19, 498–502. https://doi.org/10.1016/J.CUB.2009.01.066

Theunissen, F.E., Elie, J.E., 2014. Neural processing of natural sounds. Nat. Rev. Neurosci. 15, 355–366. https://doi.org/10.1038/nrn3731

Youn, H., Sutton, L., Smith, E., Moore, C., Wilkins, J.F., Maddieson, I., Croft, W., Bhattacharya, T., 2016. On the universal structure of human lexical semantics. Proc. Natl. Acad. Sci. 113, 1766–1771. https://doi.org/10.1073/pnas.1520752113

Zatorre, R.J., Bouffard, M., Belin, P., 2004. Sensitivity to auditory object features in human temporal neocortex. J. Neurosci. 24, 3637–3642. https://doi.org/10.1523/jneurosci.5458-03.2004

Zhao, S., Wai Yum, N., Benjamin, L., Benhamou, E., Yoneya, M., Furukawa, S., Dick, F., Slaney, M., Chait, M., 2019. Rapid ocular responses are modulated by bottom-up driven auditory salience. J. Neurosci. 39, 7703–7714. https://doi.org/10.1523/JNEUROSCI.0776-19.2019

Zoefel, B., Archer-Boyd, A., Davis, M.H., 2018. Phase entrainment of brain oscillations causally modulates neural responses to intelligible speech. Curr. Biol. 28, 401–408. https://doi.org/10.1016/J.CUB.2017.11.071

Zuk, N.J., Carney, L.H., Lalor, E.C., 2018. Preferred tempo and low-audio-frequency bias emerge from simulated sub-cortical processing of sounds with a musical beat. Front. Neurosci. 12. https://doi.org/10.3389/fnins.2018.00349

